# Molecular Epidemiology and Microbiological Characteristics of *Cryptococcus gattii* VGII isolates from China

**DOI:** 10.1101/2021.12.13.472347

**Authors:** Xuelei Zang, weixin ke, Lifeng Wang, Hua Wu, Yemei Huang, Hengyu Deng, Meng Zhou, Ningxin Wu, Dingxia Shen, Xinying Xue

## Abstract

*Cryptococcus gattii* (*C. gattii*) is a fungal pathogen that once caused an outbreak of cryptococcosis on Vancouver Island, and had spread worldwide, while few data were available in China. In this study, seven clinical isolates of *C. gattii* VGII were collected from 19 hospitals, Multi-locus Sequence Typing (MLST) analysis and whole-genome sequencing (WGS) was performed, and combined with published data for phylogenetic analysis. In addition, in vitro antifungal susceptibility testing, phenotypic analysis, and in vivo virulence studies were performed, subsequently, histopathological analysis of lung tissue was performed. *C.gattii* VGII infected patients were mainly immunocompetent male, and most of them had symptoms of central nervous system (CNS) involvement. MLST results showed that isolates from china exhibited high genetic diversity, and sequence type (ST) 7 was the major ST among the isolates. Some clinical isolates showed a close phylogenetic relationship with strains from Australia and South America. All clinical isolates did not show resistance to antifungal drugs. In addition, there was no correlation between virulence factors (temperature, melanin production, and capsule size) and virulence while in vivo experiments showed significant differences in virulence among strains. Lung fungal burden and damage to lung tissue correlated with virulence, and degree of damage to lung tissue in mice may highlight differences in virulence. Our work seeks to provide useful data for molecular epidemiology, antifungal susceptibility, and virulence differences of *C. gattii* VGII in China.

**Author Summary:** *Cryptococcus gattii* is one of the life-threatening fungal pathogens that can infect immunocompetent individuals. *C.gattii* can be invaded through the respiratory tract and can be spread to the brain, causing meningoencephalitis and even death. In 1999, an outbreak of *C. gattii* VGII occurred on Vancouver Island in Canada and spread in the Pacific Northwest region, which attracted more attention. Compared with other nations, the data of *C gattii* VGII in China was limited. To better understand the molecular epidemiology and microbiological characteristics of *C.gattii* VGII isolates from China, Bioinformatics analysis and in vivo and in vitro experiments were performed for clinical strains. Multi-locus Sequence Typing and whole-genome sequencing reveal the genetic diversity of the isolates and genetic relationships with isolates from other countries. We also investigated the antifungal drug susceptibility and virulence differences of the isolates. The authors expect that our work can provide useful data for the prevention and control of *C. gattii* VGII in China and to draw attention to *C. gattii*.

## Introduction

*C. gattii* is a specific pathogenic fungus that can infect immunocompetent individuals [1,2]. It can not only invade the respiratory tract and cause pulmonary disease but can also spread to other organs, especially the CNS, causing meningitis and even death [3,4]. In 1999, an outbreak on Vancouver Island was caused by *C. gattii* and spread to the Pacific Northwest (PNW) of the United States [5,6]. Subsequently, *C. gattii* infections had been reported in six continents.

Using the International Society for Human and Animal Mycology consensus MLST scheme, *C. gattii* was currently classified into four distinct subtypes: VGI, VGII, VGIII, and VGIV [7]. MLST revealed that the outbreak was mainly caused by the VGII genotype (including the major clonal subgroup of VGIIa and the minor clonal subgroup of VGIIb) [6]. The WGS revealed that Australia was the origin of the VGIIb subtype while South America was the origin of the VGIIa subtype[8]. The genetic variety of the *C. gattii* VGII showed the difference in virulence, and also their clinical manifestations vary with the virulence. Current knowledge supports that the virulence of VGIIa is stronger than that of VGIIb [9]. We found that VGIIb is more infectious in male patients than VGIIa [5]. Moreover, VGIIa was more likely to infect the lungs, while VGIIb was more likely to infect the CNS [6].

*C.gattii* has three classic virulence factors: temperature, capsule, and melanin. The ability to adapt and survive at 37°C is crucial for *C.gattii* to infect the host[10]. Its capsule not only inhibits phagocytosis by host phagocytes but also inhibits dendritic cell maturation and disguises antigen recognition [11]. Melanin is a reactive oxygen species (ROS) scavenger, which protects fungal pathogens from host immune responses [12].

In 2008, the first case of C. gattii VGII infection was reported in China [13], followed by other provinces [14-19]. However, these data were from a single-center retrospective study with no systematic studies on *C. gattii* VGII in China. For this purpose, *C.gattii* VGII was collected from 19 hospitals and analyzed the genetic relationships among strains. In vitro and in vivo experiments were performed to characterize the differences in virulence of *C. gattii* VGII isolates. Finally, possible virulence factors were screened to explain the differences in virulence among isolates.

## Methods

### Ethics statement

The study was conducted according to the guidelines of the Declaration of Helsinki, and approved by the Ethics Committee of Beijing Shijitan Hospital (protocol code sjtky11-1x-2020 (20)).

### Clinical isolates and patients

A total of eight strains of *C. gattii* were collected, of which five isolates have been identified as *C. gattii* VGI while three as *C. gattii* VGII by molecular typing of URA5-RFLP analysis [17]. The clinical data of patients infected by eight clinical isolates were collected. The demographic data, immune status, clinical symptoms, and other details were recorded.

### DNA extraction and sequencing for isolates

Breaking the cell wall of *C.gattii* using liquid nitrogen grinding, and the DNA of all isolates was extracted by CTAB (cetyl trimethyl ammonium bromide) method [21]. DNA of isolates was sequenced by Illumina HiSeq X Ten platform by 150 bp paired-end sequencing.

### MLST phylogenetic analysis

As described in the previous studies [22], seven unlinked genetic loci (CAP59, GPD1, IGS1, LAC1, PLB1, SOD1, and URA5) were performed independently amplified and sequenced. The STs of the clinical isolates were identified were determined by comparing the sequences with the MLST database (http://mlst.mycologylab.org/), and reference MLST sequences were also downloaded from the MLST database. In addition, we searched all related publications in PubMed to collect another 16 strains of *C. gattii* VGII information reported in China. Finally, a maximum likelihood tree was constructed based on all sequences using RAxML 8.2.12 [23].

### Detection of Single nucleotide polymorphism (SNPs) variant and analysis of WGS phylogenetic

The 7 clinical isolates were performed by WGS, and 117 additional WGS data were collected from BioProject PRJNA244927. Each Illumina data was aligned to VGII reference genome R265 by bwa-mem2 [24], and removed adapters by fastp 0.20.0 [25]. To call variants, the Picard tools (http://picard.sourceforge.net) AddOrReplaceReadGroups and MarkDuplicates were used for pre-processing, followed by GATK HaplotypeCaller 4.1.8.1 [26]. SNPs were retained if they were homozygous in all samples and passed GATK recommended standard of hard-filtering, and then were used to construct a phylogenetic tree by RAxML 8.2.12 [23] with 1000 bootstrap replications, and the final phylogenetic tree was visualized by iTOL 6 [27].

### Antifungal susceptibility testing

The Sensititre YeastOne broth microdilution system (Thermo Fisher Scientific, USA) was used to perform the antifungal susceptibility testing for six antifungal drugs, fluconazole (FCZ), voriconazole (VCZ), itraconazole (ICZ), Posaconazole (POS), amphotericin B (AMB) and 5-flucytosine(5-FC). Follow the instructions to perform antifungal sensitivity testing. Since there were no clinical breakpoints (CBPs) for the *C.gattii*, we used epidemiological cutoff values (ECVs) as reference values for VGII, FCZ: 32mg/L; VCZ: 0.25mg/L; ICZ: 0.5mg/L; POS: 0.5mg/L; AMB: 1mg/L; 5-FC: 8mg/L.

### Toxicity-related phenotypic assays

All clinical isolates were placed in yeast extract-peptone dextrose (YPD) medium and cultured overnight at 30°C. The cells were washed twice with sterile saline, centrifuged at 4000 rpm for 2 min, and the supernatant was discarded. The strains were uniformly adjusted to 5×10^7^cells/ml by counting cells on a hemocytometer, gradient diluted to 10^7^cells/ml, 2×10^6^cells/ml, 4×10^5^cells/ml, 8×10^4^cells/ml, and 1.6×10^4^cells/ml, and then performed the following assays.

#### Temperature

The cells (3 µl from each concentration) were respectively plated onto agar-based YPD medium, and cultured at 30°C, 37°C, and 39°C for three days to observe the growth at distinct temperatures.

#### Melanin production

For the melanin production assay, induction of melanin using minimal medium containing L-3,4-dihydroxyphenylalanine (L-DOPA). The cells (3µl from each concentration) were respectively cultured on a minimal medium containing L-DOPA at 30°C for three days, and photographed for melanin production [12].

#### Capsule size

Using the method described in the previous article to measure capsule diameter of isolate strains [28], cells were incubated at 30°C, stained with India ink, and acquired images using Carl Zeiss Microscopy. Finally, capsule diameter was measured manually using the cell measurement software (Zen 2011).

### In vivo murine inhalation model

All clinical isolates and reference strains (R265 and R272) were selected for virulence assay. The strains were cultured overnight at 37°C in YPD medium, after centrifugation, washed twice with sterile saline, and the final concentration of the cells was adjusted to 2×10^6^cell/ml. Female C57BL/6 mice aged 8-10 weeks were used and anesthetized with 10% chloral hydrate, 50μl fungal suspension was slowly dripped into the nasal cavity of each mouse [29].

Ten mice in each group for survival test. Observe the death of mice every day and continuously for 90 days, recorded the death of mice.

Five mice in each group for the fungal burden test, the mice were euthanized 14 days after infection. After dissection, the lungs and brains were separately placed in a 15 ml centrifuge tube containing 4 ml PBS. The tissue was smashed and homogenized in a biosafety cabinet, and then tissue suspension was serially diluted (10 times) and spread on the YPD plate. Culture at 30°C and then count Colony-Forming Unit (CFU) after two days.

Four mice in each group for pathological studies, mice infected with different strains were euthanized at 14 days and also at the time of signs of poor health. All surviving mice were euthanized at the end of the experiment (90 days). After dissection, lung tissue was fixed in formalin, embedded in paraffin, and sectioned, stained sections with Hematoxylin solution & Eosin dye (HE), then dehydrated and sealed with neutral gum. Finally, observe with microscope inspection, image acquisition, and analysis.

### Statistical Analysis

All data were analyzed using SPSS software (version 19.0; IBM Corp., Armonk, NY USA), and a descriptive statistical method was applied to summarize the clinical characteristics of patients. Continuous variables between two groups by the Manne Whitney U test. A p-value of 0.05 was considered significant. Using GraphPad Prism version 8.0 (La Jolla, CA, USA) for plotting.

### Data Availability Statement

We already have deposited the data in NCBI database. Accession to cite for these SRA data: PRJNA721774.

## Results

### Demographic and Clinical Characteristics of patients

In this study, clinical information was collected from 7 cases of *C. gattii* VGII patients (Table 1). All patients were immunocompetent, of whom six patients were male and one patient was female. The oldest patient was 83 years old, the youngest was 40 years old, and the average age was 52.7 years. All patients were diagnosed with meningoencephalitis, common clinical symptoms were headache and fever.

### MLST and WGS for phylogenetic analysis

We performed a genotypic analysis of isolates through MLST Consensus Scheme. Five STs were identified among 7 clinical isolates, including ST7 (2 strains) (as R272, the minor clone for the *C. gattii* outbreaks in Vancouver), ST21 (1 strain), ST44 (2 strains), ST173 (1 strain), and ST126 (1 strain). To better study the genetic relationships of *C. gattii* VGII in China, Sixteen strains reported so far in China were collected and combined to construct a phylogenetic tree. The result shows a high genetic variability amongst *C. gattii* VGII isolates from china, twelve STs were identified among 23 strains.ST7 was the most common ST (7/23), followed by ST44 (3/23). But no ST20 was found in China (as R265, the major clone for the *C. gattii* outbreaks in Vancouver) (Fig 1).

**Fig 1.**
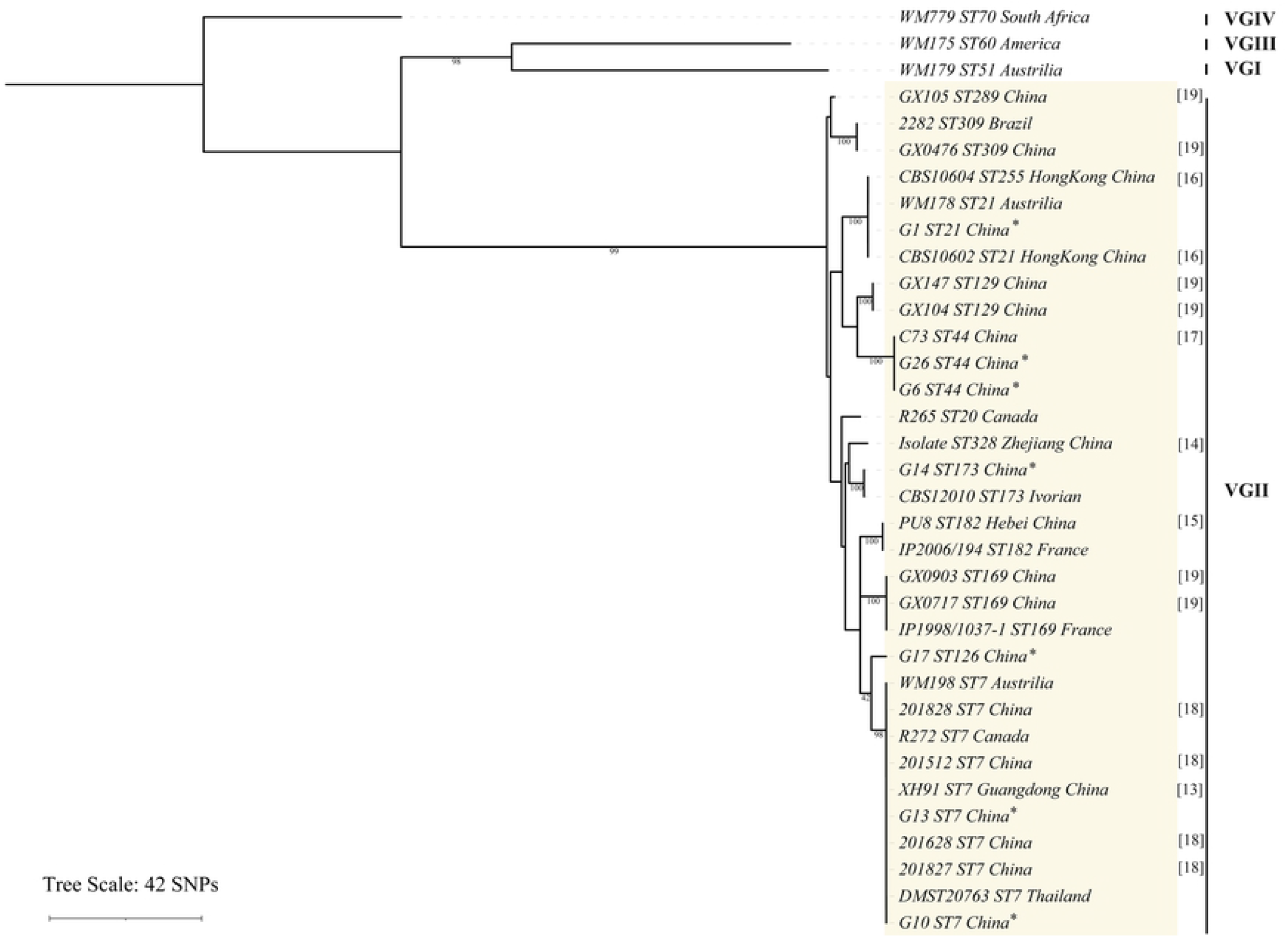
Phylogram based on MLST showing the genetic relationships between the *C.gattii* VGII isolates in the present study and another 16 isolates reported[13-19], and two clones isolate (R265, R272).

To fully understand the genetic relationships between *C. gattii* VGII strains found in China and other countries. Illumina Hi-Seq platform was used to perform the whole genome sequencing of clinical isolates, and WGS data of VGII strains published from the database was also collected. R265 was used as an outgroup to construct the phylogenetic tree. We found that the phylogenetic tree constructed based on the whole genome had similar clustering to the phylogenetic tree based on MLST. The VGII population was highly diverse, where 35492 SNPs were identified. The WGS results revealed two VGIIb strains (G10, G13) but no VGIIa strain in China. G1 showed a close genetic relationship with strains from the United States and Australia, and G14 was clustered with strains from Venezuela (Fig 2).

**Fig 2.**
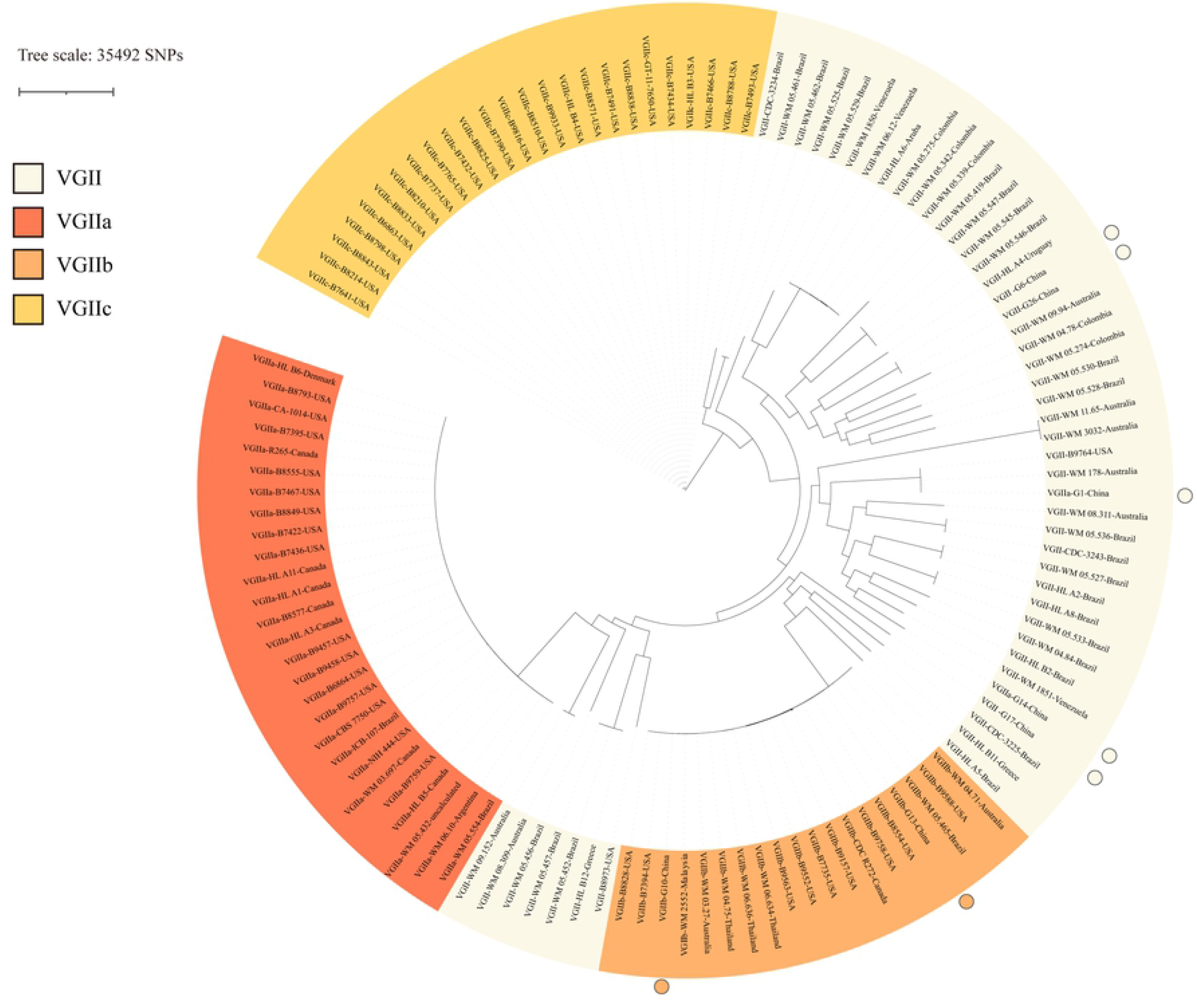
Phylogram based on WGS showing the genetic relationships between the *C.gattii* VGII isolates in the present study and global isolates from the database.

### Antifungal susceptibility testing, clinical treatment, and prognosis

All clinical isolates did not appear to be resistant to antifungal drugs, and the minimum inhibitory concentration (MIC) range was shown in Table 2. The MIC range of the strain for each antifungal drug was: 1-2mg/l for 5-FC, 0.12-0.5mg/l for POS, 0.12-0.25 mg/l for VCZ,0.12-0.25mg/l for ICZ, 0.25-4 mg/l for FCZ, 16-32 mg/l for AMB. After the diagnosis, five patients were treated with AmB+FCZ, and two patients were treated with FCZ. During the follow-ups, the patients were all alive and had a good prognosis after treatment.

### In vitro study of three virulence factors

All isolates showed similar tolerance to 37°C and showed growth well, but at 39°C, the growth of the strains exhibited remarkable differences. Compared to R265, only strain G17 was less tolerant to high temperature, while all other clinical strains showed higher resistance to high temperature (Fig 3A). The capsule sizes of the standard strains from Vancouver Island were smaller than those of the domestic clinical strains. Clinical strains had different capsule sizes, with the largest being G26 and the smallest being G6 (Fig 3B). All strains were able to be induced to produce melanin, but the production capacity of melanin was different among strains, with the lowest melanin production being G6 (Fig 3C).

**Fig 3.**
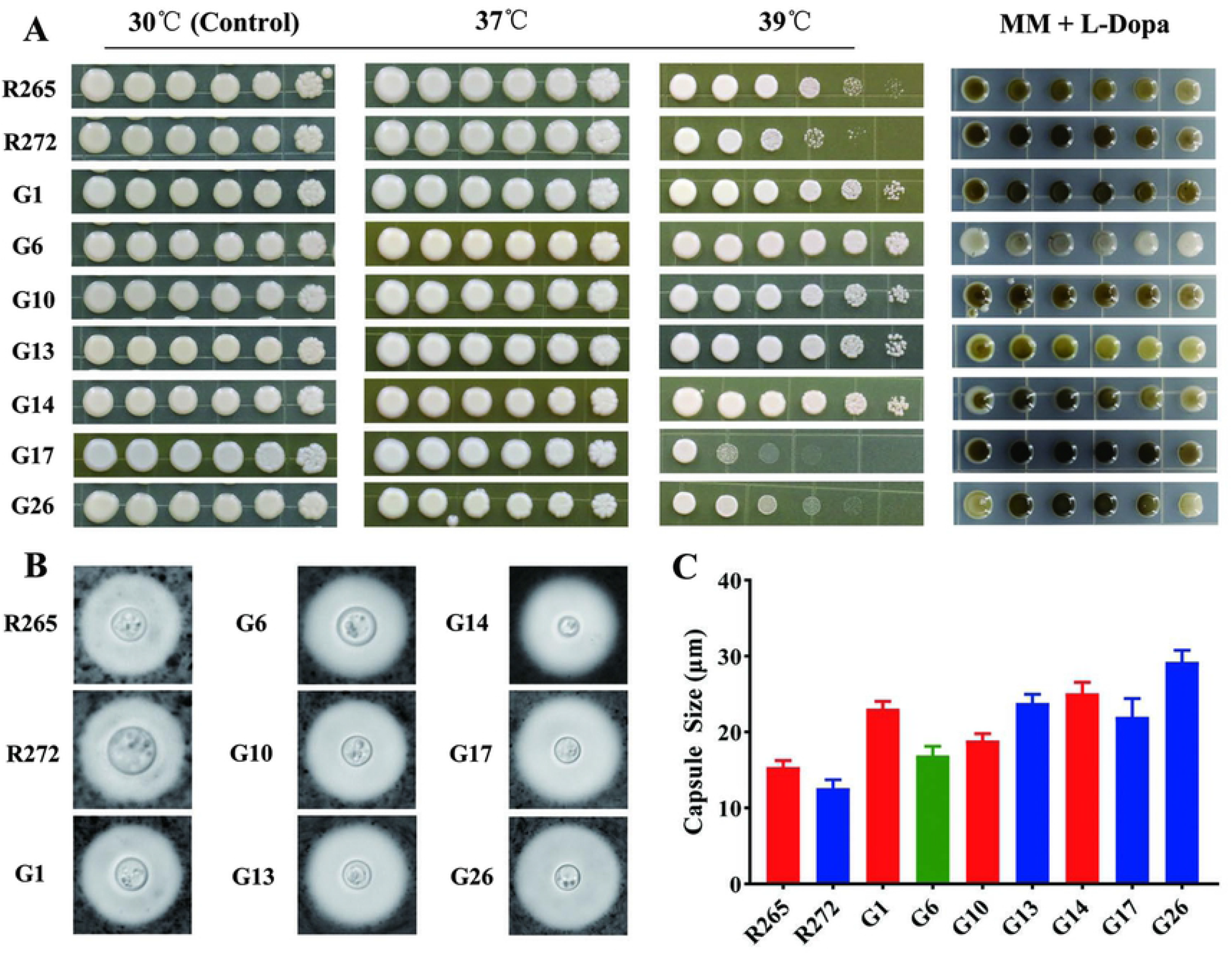
(A) Visual analysis of isolates growing in agar-based YPD medium at 37°C and 39°C for three days, and melanin production after isolates growth in minimal medium with L-DOPA at 30°C for three days. (B)Morphology of the *C.gattii* VGII isolates staining with India ink under light microscopy. (C)Capsule size was measured for 20 cells of *C.gattii* VGII isolates. Error bars indicating SD.

### In vivo virulence study in a mouse model

To evaluate the virulence difference of clinical isolates, the reference strain and all clinical strains were chosen to establish a mouse inhalation model. Fourteen days post-infection, the highest CFU counts were G14 in the lung, followed by G1 and R265, and the lowest was R272, while the highest CFU counts were G10 in the brain, followed by G17, as shown in Fig (4A, 4B).

**Fig 4.**
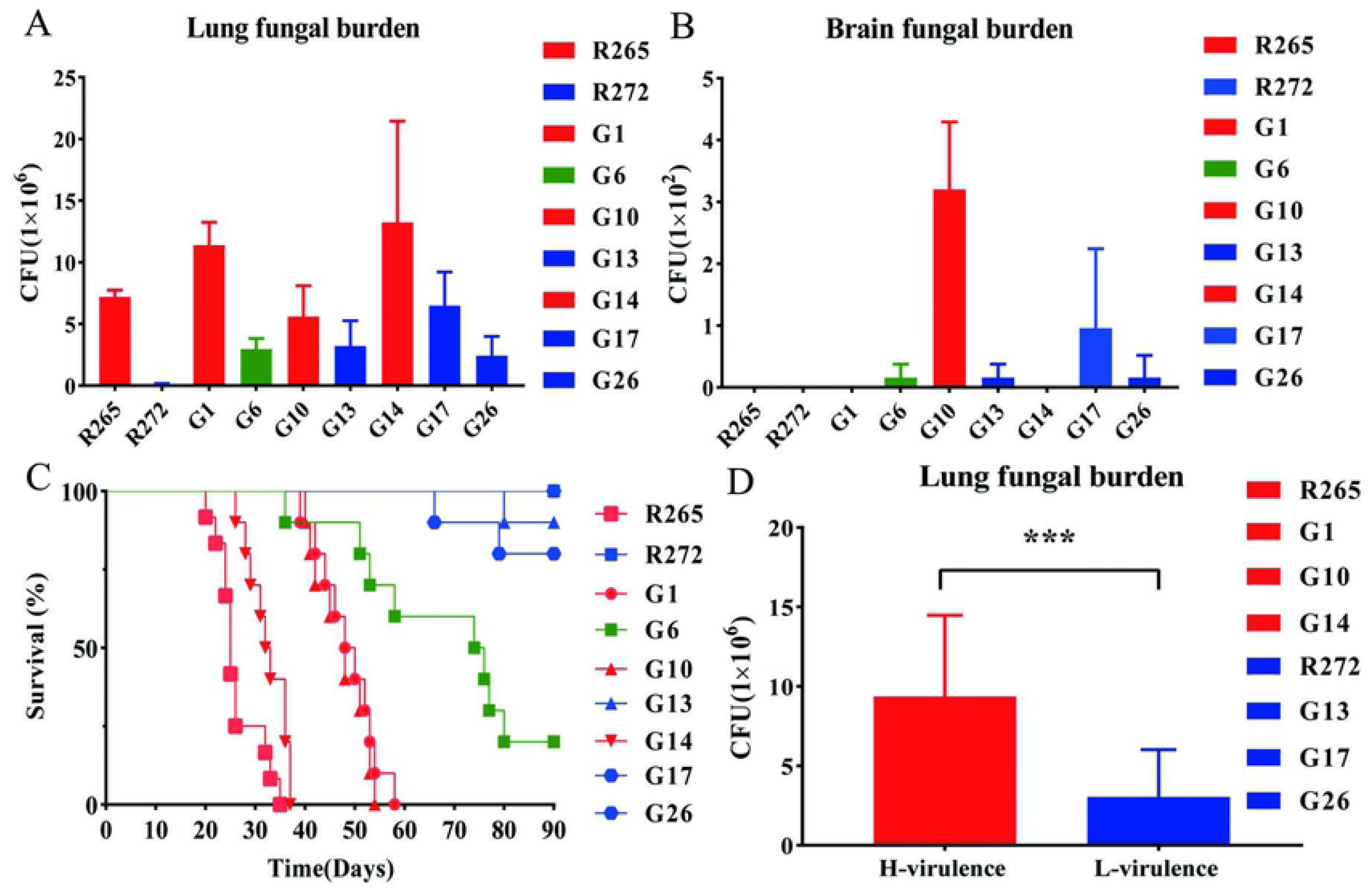
In vivo murine inhalation model. (A) (B) Fourteen days post-infection, lung and brain fungal burden of mice infected with isolates (Five mice in each isolate). (C) Survival curves of mice infected with isolates (Ten mice in each isolate). (D)The highly virulent strains (R265, G1, G10, and G14) and the less virulent strains (R272, G13, G17, and G26) were grouped and analyzed lung fungal burden difference between the two groups. Error bars indicating SD, *** < 0.001.

Survival curves showed that R265 from Vancouver Island was the most virulent strain with the minimum survival time. Each clinical strain presented a different survival time and was ordered as follows: G14, G10, G1, G6. G17 and G13 have killed one and two mice, respectively, while R272 and G26 survived until the end of the experiment, and were all considered to be less virulent strains (Fig 4C). Combined with survival curves, we found that the virulence was not significantly correlated with brain fungal burden, but an evident trend was found between virulence and lung fungal burden. Thus, the highly virulent strains (R265, G1, G10, and G14) and the less virulent strains (R272, G13, G17, and G26) were grouped and statistical analysis revealed significant differences in lung fungal burden between the two groups (Fig 4D).

Subsequently, histopathological analysis of lung tissue was performed. Interestingly, the degree of lung tissue damage was different between the high and less virulent strains. At 14 days, the less virulent strains showed normal lung tissue structure, while the high virulent strains showed the initiation of pathological injury until a late stage when obvious millet-like edema can be observed on the surface of lung tissue. H&E staining showed obvious damage and change of lung structure, and a large number of non-uniform fungal cells were dispersed in the alveolar cavity (Fig 5).We believe that the destruction of lung tissue in the mice was the primary factor causing death.

**Fig 5.**
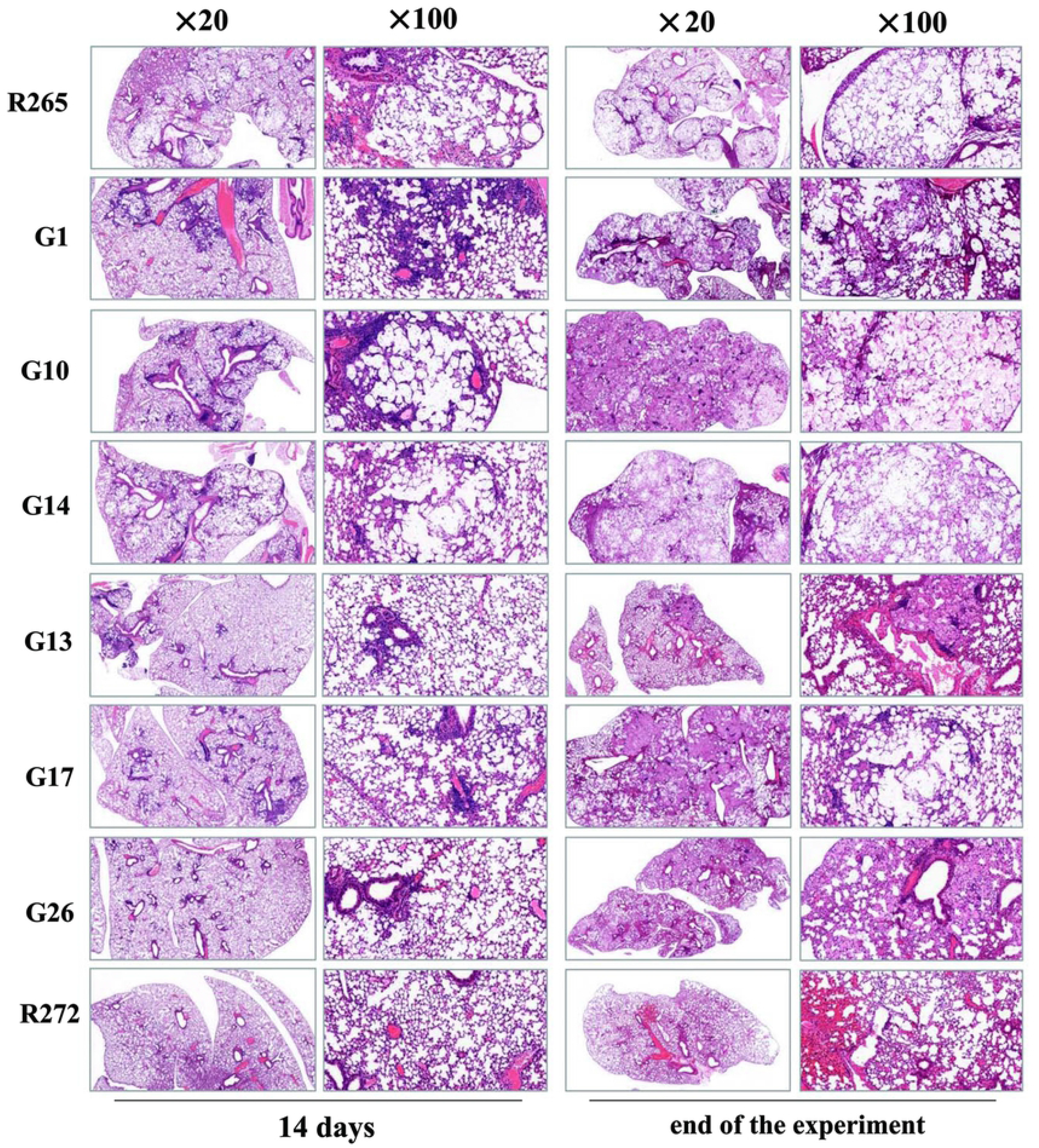
Fourteen days post-infection and end of the experiment H&E staining of the lungs of mice infected with high virulent strains and less virulent strains, the images were captured by microscope (×20, ×100), respectively.

## Discussion

In this study, we described the clinical characteristics of *C. gattii* VGII in China. In line with previous reports [30], *C. gattii* VGII is more susceptible to infecting males and immunocompetent patients. In this study, most patients had symptoms of CNS involvement, unlike previous PNW reports [5,31], most patients had symptoms of pneumonia.

A high genetic diversity amongst *C. gattii* VGII isolates in the current study, with 6 different STs. The most common ST was ST7, which was widely distributed in other parts of the world and had been reported in Canada, Brazil, Thailand, and other places [6,29,32]. According to the WGS, Australia may be the source of the outbreak on Vancouver Island, and this clonal strain spread through the eucalyptus tree to other parts of the world [33], while G10 and G13 were closely related to clonal strain (R272) from Australia, China may be the place for spread of the outbreak strain. WGS Phylogenetic tree analysis also supports the genetic relationships between some isolates and South American strains, indicating that these clonal strains may originate from South America.

The azoles MICs differ among different genotypes, among which the isolates of *C. gattii* VGII show the highest geometric mean values [34]. Clinical isolates showed relatively high MICs to FCZ, with G6 and G26 reaching 32 mg/l, which was relatively rare in the world [34]. Interestingly, both G6 and G26 are of ST44, which also provides new evidence for the different antifungal susceptibility patterns of different STs.

In vivo experiments, R265 from Vancouver Island was the most virulent strain, which was stronger than R272 [8,9] and other clinical isolates. Combined with pathology, we found that the degree of damage to lung tissue in mice was an important sign showing differences in virulence, with stronger damage to lung tissue showing greater virulence of the strains. The destruction of lung tissue was able to significantly alter the alveolar gas exchange properties, thereby causing mortality in the mice, similar to previous studies [35]. In addition, we also observed that the ability of the strains to proliferate in the lung correlated with virulence, possibly due to the invasion and massive proliferation of fungal cells in the alveolar cavity, causing the destruction of the lung tissue structure. It is also possible that fungal cell proliferation could produce some harmful substances to damage lung tissues, and the mechanism needs to be further investigated.

We performed the traditional three major virulence factor assays on all strains and found that all clinical isolates were able to grow well at 37°C, which showed the ability of the strains to consistently infect animals. However, the ability to grow at 39°C did not correlate with differences in virulence, and also the results showed that melanin production capacity and capsule size did not show a correlation with virulence either, similar to previous studies [11]. This also suggests that virulence was a very complex process and that a single phenotype among strains could hardly fully reflect the differences in virulence of strains.

Nonetheless, current strains are limited to hardly fully reflect the epidemic situation of *C.gattii* VGII in China, which is largely due to the lack of consciousness and conditions in many laboratories to identify and isolate Cryptococcus [36]. In summary, this study systematically investigates the genetic relationships, antifungal drug susceptibility, and virulence differences of C. *gattii* VGII and provides useful data for the prevention and control of *C. gattii* VGII in China.

## Author Contributions

Conceptualization: Dingxia shen. and Xinying Xue.

Methodology: Xuelei zang, Weixin Ke, Lifeng Wang, Hua Wu.

Validation: Dingxia shen, Xinying Xue.

Formal analysis: Xuelei Zang, Lifeng Wang.

Investigation: Hua Wu, Lifeng Wang, Ningxin Wu, Hengyu Deng, Meng Zhou.

Resources: Xinying Xue.

Data curation: Weixin Ke, Lifeng Wang.

Writing—original draft preparation: Xuelei Zang.

Writing—review and editing: Xuelei Zang, Dingxia shen, Xinying Xue. Visualization: Xuelei Zang.

Supervision: Dingxia shen.

Funding acquisition: Xinying Xue.

